# Microbial grazers may aid in controlling infections caused by aquatic zoosporic fungi

**DOI:** 10.1101/2020.02.03.931857

**Authors:** Hazel N. Farthing, Jiamei Jiang, Alexandra J. Henwood, Andy Fenton, Trent Garner, David R. Daversa, Matthew C. Fisher, David J. S. Montagnes

## Abstract

Free-living eukaryotic microbes may reduce animal diseases. We evaluated the dynamics by which micrograzers (primarily protozoa) apply top-down control on the chytrid *Batrachochytrium dendrobatidis* (*Bd*) a devastating, panzootic pathogen of amphibians. Although micrograzers consumed zoospores (∼3 µm), the dispersal stage of chytrids, not all species grew monoxenically on zoospores. However, the ubiquitous ciliate *Tetrahymena pyriformis*, which likely co-occurs with *Bd*, grew at near its maximum rate (*r* = 1.7 d^-1^). A functional response (ingestion vs. prey abundance) for *T. pyriformis*, measured using spore-surrogates (microspheres) revealed maximum ingestion (*I*_max_) of 1.63 × 10^3^ zoospores d^-1^, with a half saturation constant (*k*) of 5.75 × 10^3^ zoospores ml^-1^. Using these growth and grazing data we developed and assessed a population model that incorporated chytrid-host and micrograzer dynamics. Simulations using our data and realistic parameters obtained from the literature suggested that micrograzers could control *Bd* and potentially prevent chytridiomycosis (defined as 10^4^ sporangia host^-1^). However, simulated inferior micrograzers (0.7 x *I*_max_ and 1.5 x *k*) did not prevent chytridiomycosis, although they ultimately reduced pathogen abundance to below levels resulting in disease. These findings indicate how micrograzer responses can be applied when modelling disease dynamics for *Bd* and other zoosporic fungi.

## Introduction

Although the traditional microbial food web (i.e. prokaryotes and protists, *sensu* Azam et al. [1] is well-established as a driver of aquatic productivity [2], micro-fungi are only now being appreciated as integral aquatic microbes. A dominant group of micro-fungi, the chytrids, are parasites of phytoplankton, zooplankton, and vertebrates [3], and zoospores, the chytrid dispersal stage, are argued to be nutritious and of an appropriate size for protozoan grazers [2, 3]. Hence, through top-down control micrograzers within the microbial food web have the potential to reduce the likelihood or severity of, or even prevent, disease outbreaks caused by these pathogens [3-5]. Here, by developing and parameterizing a population model we explore the dynamics by which microbial grazers may control the chytrid *Batrachochytrium dendrobatidis*, a panzootic pathogen of amphibians that is argued to have caused the greatest loss of biodiversity attributed to any disease [6].

*Batrachochytrium dendrobatidis* (henceforth, *Bd*) infects amphibian hosts through the dispersal of motile, 3-5 µm zoospores (Fig. 1). The environmental pool of zoospores is instrumental in driving infection dynamics, as these can accrue in a dose-dependent manner [7], with for some hosts the size of the zoospore pool influencing long-term consequences for population survival or extinction [8]. It follows that processes that reduce the zoospore-pool should reduce the probability and intensity of disease outbreaks. Consumption of zoospores by naturally occurring micrograzers has been suggested to result in losses sufficient to reduce infections. Experiments show that some micrograzers may reduce the likelihood of *Bd* infections, and field data indicate a negative relationship between potential-grazer abundance and both the prevalence of infection and host mortality from disease [9-12]. There is now a need to build on these observations and investigate in more depth the dynamics by which consumers may impact on *Bd*.

**Fig. 1.**
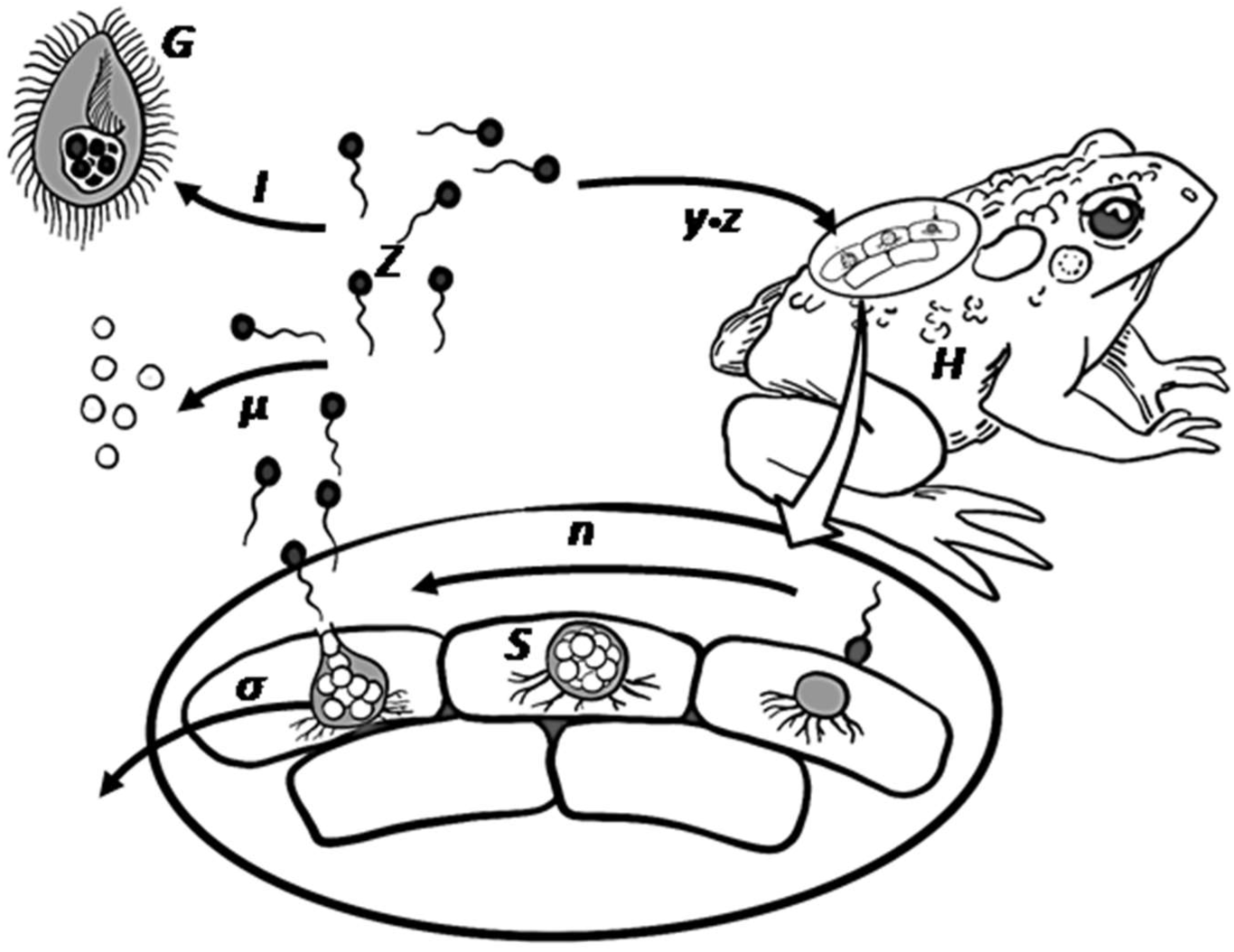
*Bd* infectious life cycle including the potential grazing pressure by micrograzers. Zoospores (*Z*) move using a flagellum, and on contact infect the amphibian host. Within the host epidermal cells, a spore then forms a sporangium (*S*) that releases further zoospores through asexual reproduction (*n*), after which the sporangium dies (*σ*). Released zoospores may die naturally (*µ*) or be ingested (*I*) by micrograzers (*G*).

Most work to date on the consumption of *Bd* zoospores has focused on large zooplankton, especially cladocerans [10, 13-15]. However, experiments on cladocerans have used unrealistically high micrograzer abundances (>10-100 times higher than natural levels), and at natural levels large zooplankton seem to have little impact on *Bd* infections [16]. Micrograzers, in contrast, are abundant in shallow waters and are often near the bottom of ponds where infected hosts (e.g. benthic, grazing tadpoles) spend time resting and grazing on the substrate [17-19]. Furthermore, as many protozoa have generation times on the order of hours, by reproducing asexually when zoospores are abundant, micrograzer populations may increase several fold, consuming zoospores as they are released from the host. We, therefore, argue that protozoa will be more important than cladocerans in reducing the abundance of chytrid zoospores. This is supported by Schmeller et al. [12] who, using mesocosms, showed the ciliate *Paramecium* can significantly reduce the number of hosts infected with *Bd* by up to 65% when it is introduced at naturally occurring abundances.

We also suggest that the main impact of micrograzers on *Bd* spore-load will be in the water directly surrounding the host, where zoospores will be most abundant. Field studies suggest that in water bodies where *Bd* occurs, zoospore densities in the water column are low, ranging from ∼0.5 to 500 L^−1^ [20, 21]. In contrast, zoospores are shed from hosts at up to 250 zoospores min^-1^ [22], surviving only ∼1-3 days [9]. Additionally, zoospores mostly disperse on the order of 1 cm [23], demonstrating chemotaxis towards keratinised skin over this distance [24, 25]. Although these laboratory-based rates will be dependent on environmental factors such as temperature [9], they suggest that the limited movement and survival of the rapidly produced zoospores will lead to dense aggregations in localized regions around the host. Recognising the likelihood of these local abundances and the well-established density-dependent feeding and growth responses of micrograzers [2], in this study we focused attention on the impact of micrograzers on *Bd* dynamics around the host. We achieved this by: first, investigating a range of potential micrograzers, determining which survived on a diet of only *Bd* zoospores, and concentrating on those that grew; second, measuring ingestion and growth rates of a common species that thrives on *Bd*; and third, using these data, developing and exploring a model that couples the *Bd* life-cycle with micrograzer-control on zoospores. In doing so, we indicate the dynamics by which micrograzers may reduce *Bd* populations – potentially preventing disease – and provide a mechanism by which chytrid-diseases can be incorporated into microbial food web models.

## Materials and methods

### Culture maintenance

*Batrachochytrium dendrobatidis* (*Bd*) cultures (strain #262 IA 9’13, Imperial Collage London) were maintained at 18 °C (at which all experiments were conducted) on H-broth medium (500 mL: 5 g Tryptone and 16 g glucose) and were regularly transferred (every ∼5 d) to maintain exponential growth. Bacterial growth was prevented by the addition of antibiotics (Ampicillin at 100 µg ml^-1^; Kanamycin at 50 µg ml^-1^; Chloramphenicol at 34 µg ml^-1^). Micrograzers were obtained from Sciento (Manchester, UK): the ciliates *Blepharisma* sp., *Oxytrichia* sp., *Paramecium aurelia, Paramecium caudatum, Stentor coeruleus, Tetrahymena pyriformis*, and *Urocentrum turbo* and the rotifers *Brachionus calcyciflorous* and *Philodina* sp. *Tetrahymena pyriformis* was maintained axenically for extended periods on H-broth. All other species were maintained prior to experiments on a natural assemblage of bacteria in Chalkley’s medium enriched with cereal grains, as provided by Sciento [26].

### *Assessing growth of micrograzer species on* Bd zoospores

We tested the hypothesis that *Bd* zoospores were of nutritional benefit to the micrograzer. To do so, we compared growth with zoospores to when no food was available. We also compared growth on zoospores to maximal rate of growth of the micrograzers, obtained from literature estimates (Fig 2).

**Fig. 2.**
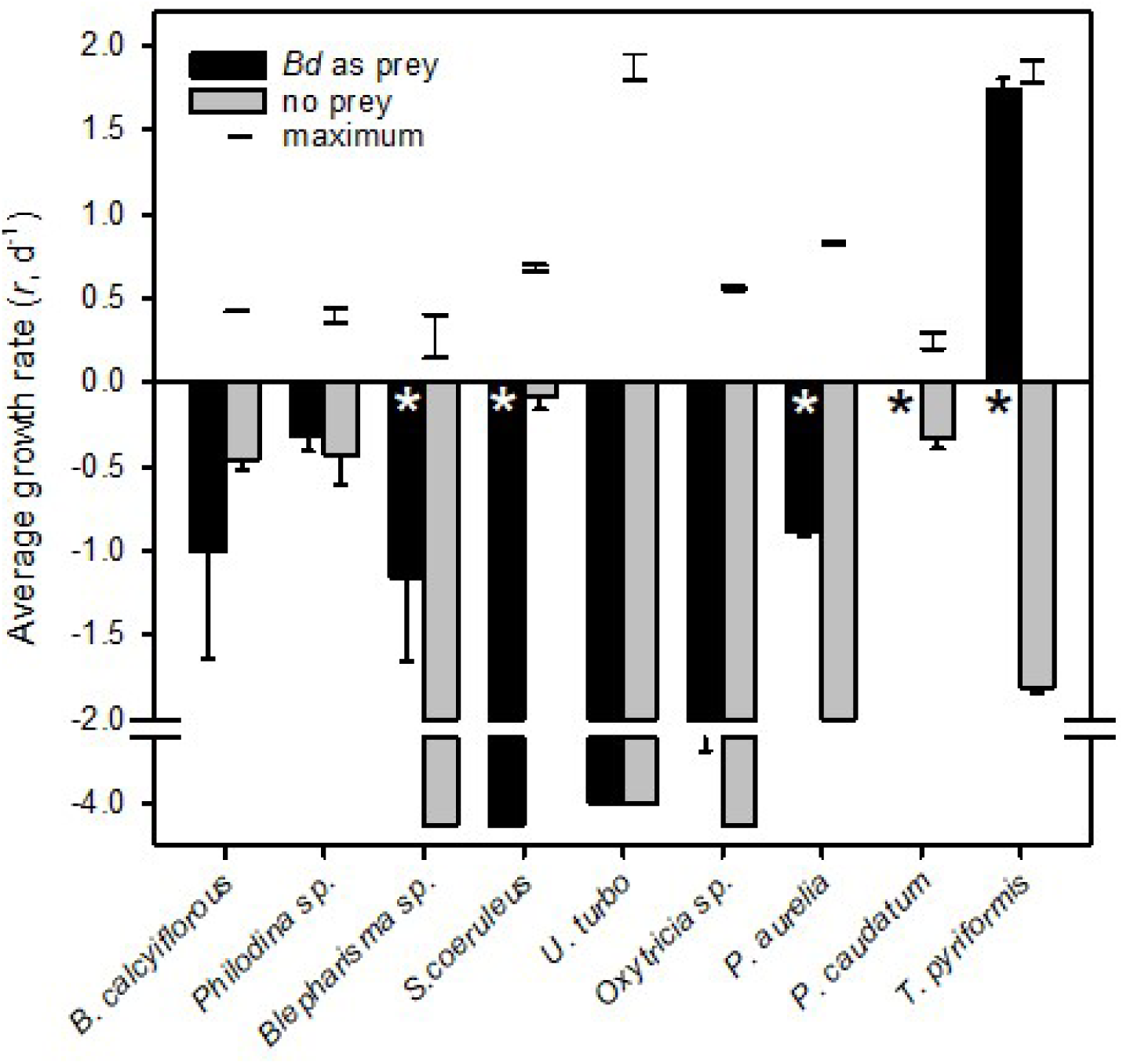
Average growth rates (*r*, d^-1^) for micrograzers fed *Bd* (black) or no prey (grey); error bars are one standard error, and * indicates where significant (α = 0.05) differences occurred between fed and unfed treatments. Note that for *P. caudatum* growth rate was zero when fed *Bd*. The horizontal lines connected by a vertical line represent the range of predicted maximum growth rate (i.e. food saturated on suitable prey) at 18 °C, for each species. Data were obtained from a various sources at a range of temperatures [59-66] and were converted to rates at 18 °C by two methods, either assuming a Q_10_ of 2 or that growth rate varies linearly with temperature at a rate of 0.07 *r* (d^-1^) °C^-1^ [67].

Prior to introducing micrograzers, *Bd* was isolated from its growth medium to ensure that the medium was not a source of nutrients for the micrograzers. To do so, a *Bd* suspension (in exponential phase) was centrifuged (50 ml tubes, 5000 rpm, 6 min), the supernatant removed, and the pellet resuspended in autoclaved Volvic^®^ mineral water to a concentration > 1.50 × 10^5^ ml^-1^ (determined microscopically). Bacterial growth was prevented with antibiotics, as above.

To assess growth rate, we followed our previous methods [27]. Micrograzers (9 species) were passed 5 times through autoclaved Volvic^®^ water to remove bacteria. Then 5 to 8 individuals, dependent on micrograzer size, were added to a 10 ml well containing the *Bd* suspension. Parallel treatments containing only sterile Volvic^®^ water were used to assess mortality rate in the absence of prey. All treatments (i.e. species incubations with or without *Bd*) were replicated in triplicate (i.e. three 10 ml wells). To assess growth rate (*r*, d^-1^), the number of gazers in each well was determined microscopically after 2 or 3 days (depending on the observed change in abundance). Then, new *Bd* suspensions were prepared (as above), and micrograzers were transferred to these, maintaining *Bd* abundance. If micrograzer numbers decreased (net death occurred) over the incubation, then all individuals were transferred, but if numbers increased (net growth occurred) then the initial number was transferred. This procedure was continued for 14 days or until all micrograzers had died. Cultures were routinely checked to ensure there was no bacterial contamination.

When numbers increased between transfers, growth rate (*r*, d^-1^) was determined over each incubation period, as *r* = *ln*(*N*_t_/*N*_0_)/*t*, where *N*_0_ and *N*_t_ are the micrograzer abundance on the initial and final day respectively, and *t* is the incubation period (2 or 3 days); to determine growth rate across all transfers (up to 14 d), the average of these was obtained. When micrograzer numbers decreased between transfers, mortality rates (-*r*, d^-1^) were determined as slope of *ln* numbers over the entire incubation period. To assess if growth (or death) rate differed between treatments (i.e. with or without *Bd*) a two tailed t-test was conducted (α = 0.05).

### *Quantifying the functional response of* Tetrahymena *grazing on* Bd

Our study focused on *Tetrahymena pyriformis* as: 1) it grew rapidly on *Bd* zoospores (see Results) and therefore clearly consumed and assimilated zoospores; 2) it is a model organism for which there are substantial data (see Discussion), and 3) it is common, globally, in habitats where *Bd* may occur (see Discussion). Prior to determining ingestion rate, *T. pyriformis* was acclimated with zoospores for >10 h. To do so, the ciliates were first removed from H-broth by centrifugation (50 ml tubes, 8000 rpm, 8 min) and then resuspended in 10 ml of autoclaved Volvic^®^ water. To obtain only zoospores, a centrifuged *Bd* culture (as above) was filtered through a 5 µm Nitex^®^ mesh. Zoospores were then added to the water containing ciliates, to a total volume of 20 ml (resulting in ∼10^6^ zoospores ml^-1^), with antibiotics (as above). This acclimation had no negative effects on the ciliates: after 10 h, zoospore abundance had substantially decreased and ciliate abundance increased (indicating the ciliates were feeding and growing), the ciliates behaved similarly to when grown on H-broth (i.e. similar swimming pattern), and their cell size and shape were similar to when grown on H-broth.

To determine ingestion rate on spore-sized particles, 3 μm fluorescent polymer microspheres (henceforth beads, Fluoro-Max(tm), Thermo Fisher Scientific, USA,) acted as a surrogate for *Bd* zoospores which are 3 - 5 µm [28]. Bead concentrations, ∼8 × 10^3^ ml^-1^ to 10^6^ ml^-1^ (see Results), were prepared in autoclaved Volvic^®^ water and vortexed prior to use, ensuring mono-dispersion. An aliquot (0.5 ml) of the acclimated ciliate culture (> 30 micrograzers) was added to 1 ml of Volvic^®^ water with beads, at various concentrations (with more measurements at low abundances [29], see Results), and incubated for 5 or 10 min, depending on the bead concentration (preliminary experiments deemed these to be appropriate incubation periods). Incubations were terminated by fixing cells with ethanol (final concertation 70%). The average number of beads ingested per ciliate (>30 cells) was determined via fluorescent microscopy and was subsequently used to determine ingestion rate (*I*, prey d^-1^) at each prey concentration.

The relationship between ingestion rate and zoospore abundance (*Z* ml^-1^), was determined by fitting a Type II functional response to the data: *I* = *I*_max_\**Z*/(*k* + *Z*), where *I*_max_ (*Z* min^-1^) is the maximum ingestion rate and *k* is the half saturation constant (*Z* ml^-1^). The response was fit using the Marquardt-Levenberg algorithm (SigmaPlot, Systat, Germany); this algorithm is appropriate for describing such biological data sets [30].

### *Modelling micrograzer impacts on* Bd *populations*

To assess the dynamics by which grazing pressure impacts on *Bd* infection populations we developed and applied the following model that couples a reduced version of the *Bd*-load model [8] with the Rosenzweig-MacArthur predator-prey model [31]. Data for *T. pyriformis* were used to represent micrograzers (see the Discussion for a justification to focus on this species). Following logic outlined in the Introduction, the model describes the infection load on a single host and, as a proxy for the waters surrounding the host, only considers a volume of 10 ml around that host, where zoospores and micrograzers reside. As a metric to predict chytridiomycosis, it assumes that a sporangia load of 10^4^ per host results in host mortality [8], with the recognition that this will vary between hosts and *Bd* strains [32, 33]. It does not include reduction of spore numbers by emigration as zoospores are unlikely to disperse far before dying [23], and we assume through chemokinesis, micrograzers remain near their food source[34-36]. The model is described by the following equations,

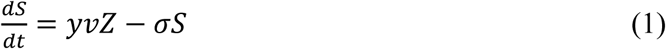

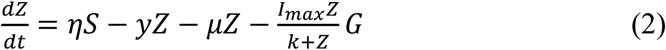

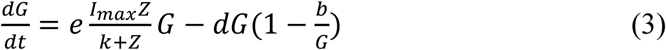

where for Eq. 1 and Eq. 2, *S* is the number of sporangia ml^-1^ (note for per host measurements this value is multiplied by 10); *Z* is the zoospore abundance (ml^-1^); *y* is the per capita spore-host encounter rate; *v* is the fractional likelihood of spore infection per encounter; *σ* is the per capita sporangia mortality rate; *η* is the per sporangia spore-release rate; and *µ* is the per capita spore mortality rate (see Table 1).

**Table 1.** Parameters used to assess *Bd* dynamics (Eq. 1-3). *Bd* parameter estimates are from Briggs et al. [8] who present a wide range of possible values; we have chosen one set of these that provide an illustration of Bd-micrograzer dynamics. Micrograzer data are from our functional response (Fig. 4). Conversion efficiency (*e*) was estimated as described in the text. The minimum number of micrograzers (*b*) and micrograzer death rate (d) were derived from personal observations.

| Symbol | Parameter | Estimate | Range explored by Briggs et al. [7] | Dimension |
| --- | --- | --- | --- | --- |
| $\gamma$ | Rate of zoospores encounter with hosts | (0.05) | Large range of values | $\text{d}^{-1}$ |
| $\nu$ | Fraction of successful *Bd* spore infections | 0.1 | 0-1 | dimensionless |
| $\eta$ | Production rate of zoospores from sporangium | 17.5 | -- | $\text{d}^{-1}$ |
| $\sigma$ | Sporangia loss rate | 0.2 | 0.1-0.3 | $\text{d}^{-1}$ |
| $\mu$ | Spore death rate | 0.1 (1) | 0.02-1 | $\text{t}^{-1}$ |
| $E$ | Conversion efficiency | $5 \times 10^{-4}$ | | dimensionless |
| $I_{\max}$ | Maximum ingestion rate | 1630 | | $\text{S d}^{-1}$ |
| $K$ | Half saturation constant | $5.75 \times 10^4$ | | $\text{S ml}^{-1}$ |
| $d$ | Micrograzer death rate | 0.01 | | $\text{t}^{-1}$ |
| $b$ | Minimum number of micrograzers | 1 | | $\text{ml}^{-1}$ |

Then, Eq. 2 and Eq. 3 were coupled to include spore loss by micrograzers (*G*), where grazing (*I*) is dictated by the functional response (see *Tetrahymena pyriformis* ingestion, above); *e* is the abundance-based conversion efficiency, determined assuming a biomass-based estimate of *e* is ∼0.1 [37] and biovolumes of *Bd* zoospores and *T. pyriformis*; and *d* is the micrograzer per capita death rate. We assume here that *Bd* zoospores are not the only potential food source for the micrograzers, and so incorporate a minimum micrograzer abundance (*b*) that implicitly assumes that in the absence of zoospores the micrograzer population is maintained by the presence of other potential food sources; hence we model potential increases in micrograzer abundance over and above this minimum, dependent on consumption of *Bd* zoospores. Estimates of *d* and *b* are based on our unpublished observations. Table 1 summarises the above parameters and the estimates used.

All model runs (100 d) were initiated with 10 sporangia host^-1^ (1 sporangium ml^-1^), 100 zoospores ml^-1^, and 1 micrograzer ml^-1^ (again assumed to be the minimal population size, maintained by other resources in the environment). For *Bd*, we applied parameter values that were within the range explored by Briggs *et al*. [8] (Table 1).

We first performed simulations to assess the ability of *T. pyriformis* to control *Bd*. Then, we assessed the extent to which micrograzers that are inferior to *T. pyriformis* could control *Bd*, through exploration of micrograzer parameter space. Inferior micrograzers had reduced maximum ingestion rate (up to 0.5 × *I*_max_ of *T. pyriformis*) and increased half saturation constant (up to 2 × *k* of *T. pyriformis*); see Fig. 4 for an indication of the range of these responses. To quantify the impact of micrograzers on *Bd*, we examined the maximum abundance (over the 100 days) and the abundances at 50 and 100 days of *S, Z*, and *G*.

## Results

### *Assessing growth of micrograzer species on* Bd

All of the micrograzers can be maintained in laboratory cultures, with maximum growth rates ranging from ∼0.4 to 2 d^-1^ (Fig 2), and all died when maintained on water alone, indicating their relative mortality rates when starved (Fig. 2). When fed *Bd*, micrograzers exhibited four distinct responses (Fig. 2): 1) for the ciliate *Stentor coeruleus* the death rate was significantly (and substantially) higher than in water alone; 2) for the ciliates *Urocentrum turbo, Blepharisma* sp., and *Oxytrichia* sp. and the rotifer *Philodina* sp. there was no significant difference between death rate with or without *Bd*; 3) for the rotifer *Brachionus calcyciflorous* growth rate initially increased (i.e. after 48 h) followed by death, and for the ciliates *Paramecium aurelia* and *P. caudatum* (Fig. 3) the growth rate was initially positive when *Bd* was present followed by a negative growth rate as time progressed – on average over the incubation *P. aurelia* exhibited negative growth while *P. caudatum* exhibited zero growth (Fig. 3); and 4) for the ciliate *Tetrahymena pyriformis* there was a sustained positive growth rate (Fig. 2), which was significantly higher than the negative growth rate on water alone; this growth rate of ∼1.7 ± 0.23 (SE) d^-1^ was equal to that when the ciliate was grown axenically on H-broth (unpublished data) and near its maximum rate under any conditions.

**Fig. 3.**
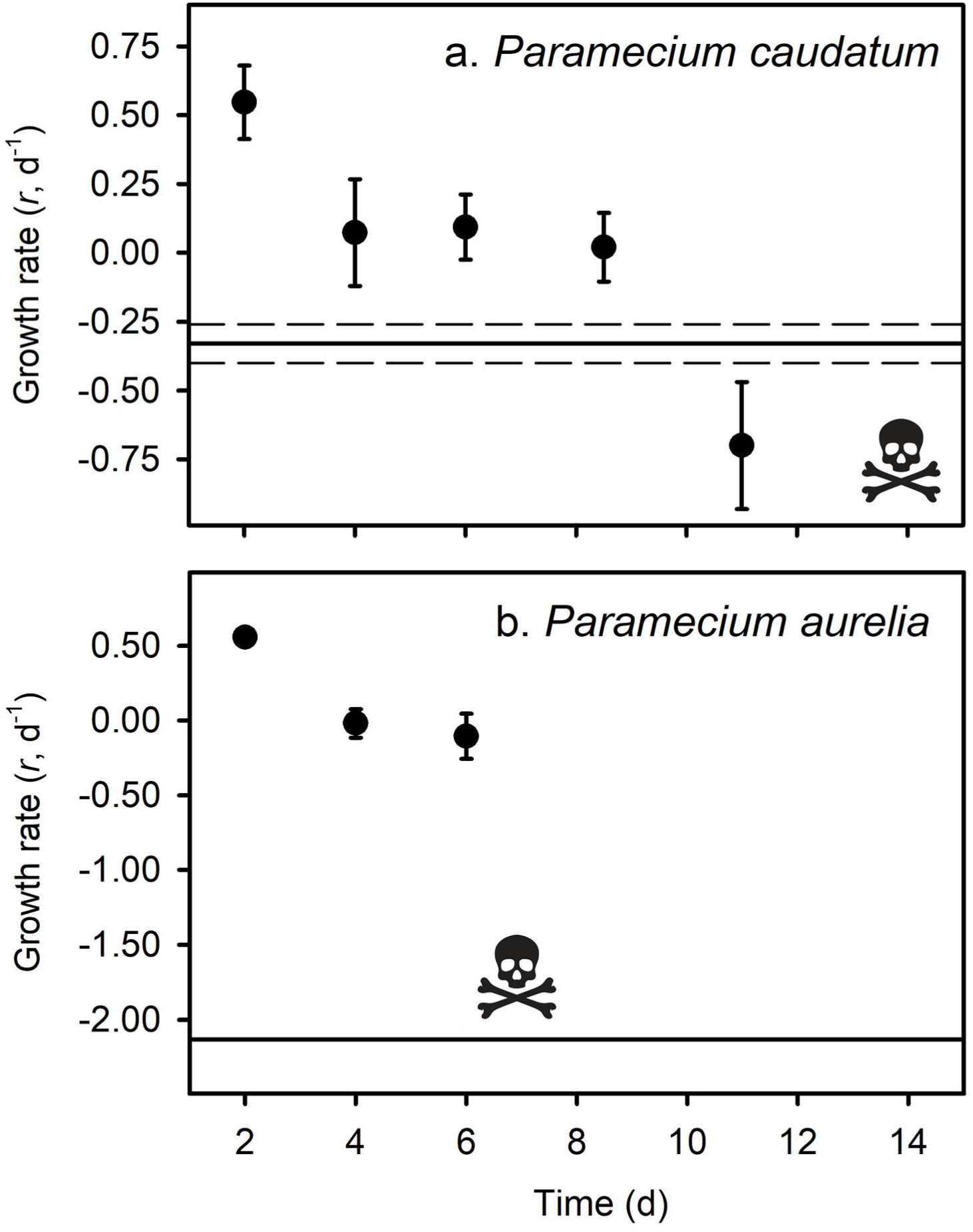
Average growth rates (*r* d^-1^) of three replicates of *Paramecium caudatum* (a) and *Paramecium aurelia* (b) in the *Bd* treatment, with standard error bars. The skull and crossbones indicate the time point where all individuals had died. The solid black line represents the average death rate of the micrograzers when no prey were present, and the dotted black line indicates the standard error of the control groups.

### *Quantifying the functional response of* Tetrahymena *grazing on* Bd

As *T. pyriformis* grew on zoospores alone it was clear that this ciliate ingested *Bd* zoospores. Zoospores were also observed in *T. pyriformis*, in the food vacuoles of the ciliate, under 40 x magnification. Ingestion rate followed a typical Type II (rectangular hyperbolic) functional response (Fig. 4, adjusted *R*^2^ = 0.82), with *I*_max_ = 1.63 × 10^3^ ± 98 (SE) prey d^-1^ and *k* = 5.75 × 10^3^ ± 1.38 × 10^3^ (SE) prey ml^-1^.

**Fig. 4.**
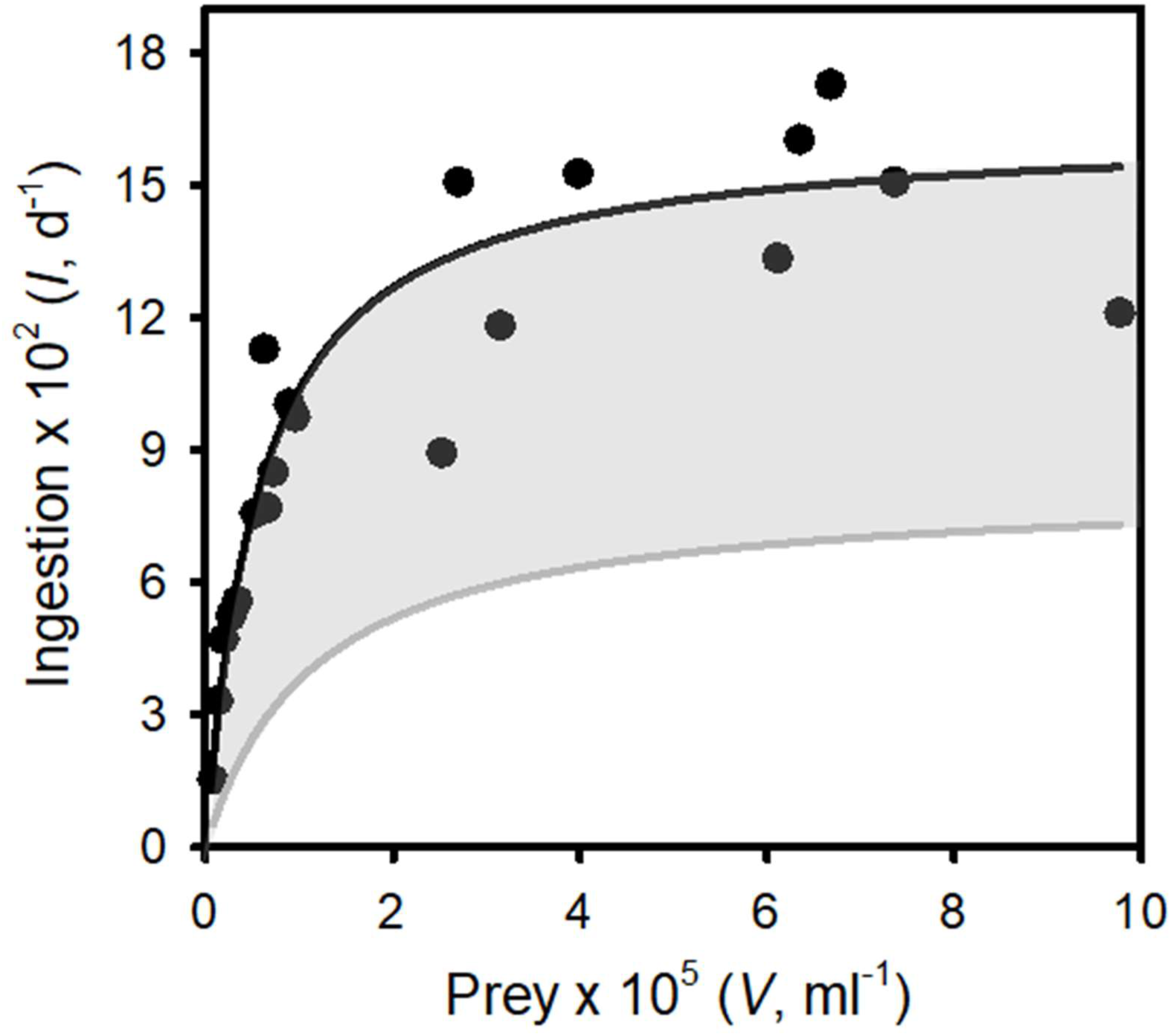
The functional response: ingestion rates of *Tetrahymena pyriformis* on surrogate zoospores (prey) vs prey concentration. Points are ingestion rates at defined prey abundances. The solid line represents the best fit of a Type II functional response to the data (see Results for the parameter estimates). The grey region represents the range of functional responses used to assess the ability of “inferior micrograzers” to control *Bd* (i.e. reduced maximum ingestion rate and increased half saturation constant; see Methods, Modelling micrograzer impacts on *Bd* populations).

### *Modelling micrograzer impacts on* Bd *populations*

Control of *Bd* occurred when parameters for the micrograzer (*T. pyriformis*) were included in the model (Fig. 5). In the absence of the micrograzer, sporangia per host reached lethal levels (>10^4^ host^-1^ [8]) by ∼55 days (Fig. 5a). However, when micrograzers were included their population rose from 1 to ∼35 ml^-1^, with the result that sporangia were limited to a maximum of 3 × 10^3^ per host (i.e. based on the assumption that 10^4^ sporangia is a lethal limit, the host would survive), and *Bd* was virtually eradicated by 100 days (Fig. 5b).

**Fig. 5.**
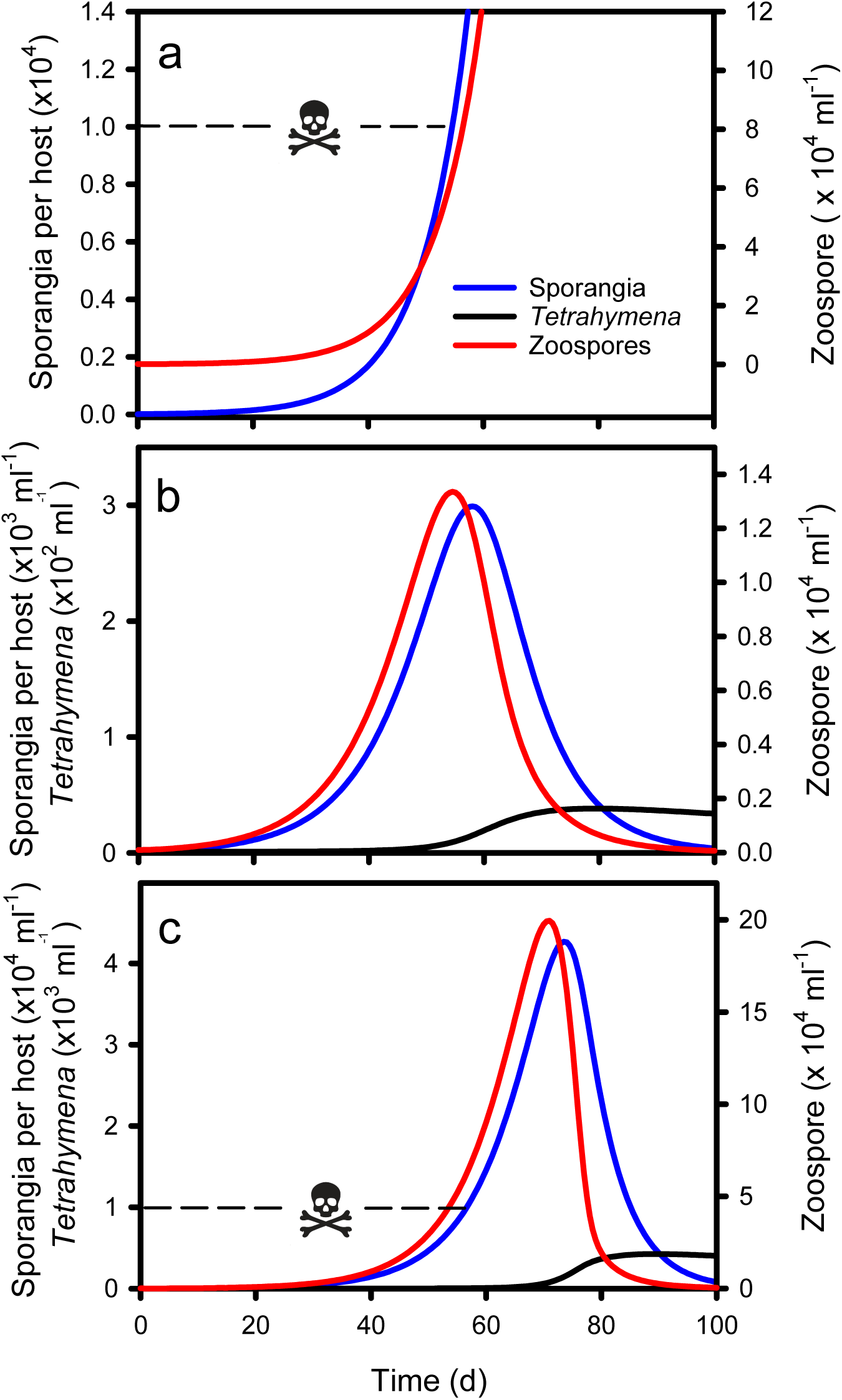
Simulations of micrograzer (*T. pyriformis*) control of *Bd*, based on Eq. 1-3 and parameters presented in Table 1. a. *Bd* (zoospore and sporangia) dynamics in the absence of micrograzers, indicating that by ∼55 days sporangia per host approach lethal limits (skull and crossbones, 10^4^ sporangia per host Briggs et al. [8]). b. *Bd* and micrograzer dynamics, indicating control of zoospores and sporangia, maintaining sporangia numbers below the lethal limit. c. *Bd* and micrograzer dynamics based on an inferior micrograzer to *T. pyriformis* (0.5 x *I*_max_; 2 x *k* presented in Table 1), indicating the micrograzers inability to prevent host death at ∼55 days (skull and crossbones) but its ability to ultimately reduce *Bd* levels by 100 d.

We then assessed the ability of micrograzers that were inferior to *T. pyriformis* to control *Bd*, through exploration of micrograzer parameter space: i.e. up to twice the half saturation (*k*) and half the maximum ingestion rate (*I*_max_) of *T. pyriformis* (Fig. 4). Fig. 5c illustrates population dynamics when the most inferior micrograzer was included (highest half-saturation constant and lowest maximum ingestion rate): the general pattern remained similar to that when *T. pyriformis* parameters were applied, with the micrograzers controlling *Bd* over 100 d, but the abundance of zoospores, sporangia, and micrograzers were more than 10 times greater than the simulation including *T. pyriformis*, leading to predicted host death at ∼55 days and a peak in abundance at ∼70 days.

We then examined the pattern of the temporal dynamics across a wider range of parameter space (representing a range of predators-types) by reporting the maximum abundance and the abundances at 50 and 100 days of zoospores, sporangia, and micrograzers. Across all parameters explored, the micrograzer population provided top-down control of *Bd*, as over the entire range *Bd* was virtually eradicated by 100 days (Fig. 6 c, f). However, the quantitative levels and rates of control varied considerably with micrograzer efficiency: with reduced *I*_max_ and increased *k*, zoospores and sporangia reached higher maximum abundances (Fig. 6a, d) and persisted longer (Fig. 6b,e), indicating a decrease in the control of *Bd*. In particular, micrograzers with < 0.7 *I*_max_ (∼10^3^ prey d^-1^) and >1.5 x *k* (∼9 × 10^4^ ml^-1^) were not capable of preventing sporangia per host exceeding lethal levels of 10^4^ per host (the yellow-to-red region on Fig. 6 d). Decreased *I*_max_ and increased *k* also led to increases in micrograzer abundance (Fig 6. g-i), in response to the increased spore levels available under these grazing regimes.

**Fig. 6.**
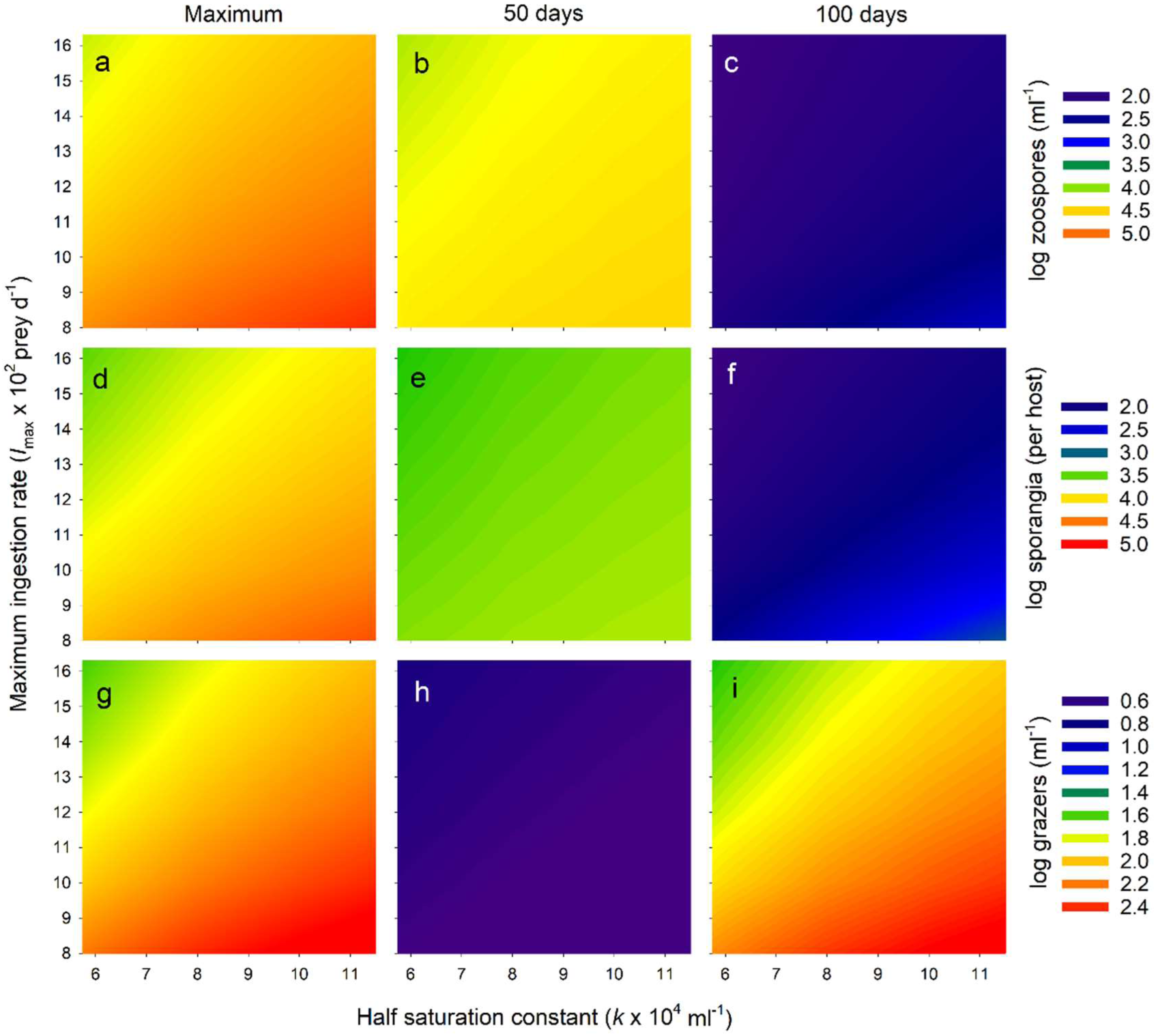
Exploration of *Bd*-micrograzer dynamics (Eq. 1-3), through varying two key micrograzer parameters: the half saturation constant (*k*) and the maximum ingestion rate (*I*_max_); see Methods for details. To characterise dynamics, we provide the log numbers of zoospores, sporangia, and micrograzers. For each, we present the maximum number reached over the 100 days, the number at 50 days, and the number at 100 days.

## Discussion

Control of a wide range of diseases caused by zoosporic fungi may be achieved through consumption on aquatic, motile zoospores by micrograzers [3-5]. Here, we explore the dynamics by which micrograzers may play a pivotal role in controlling the devastating amphibian disease chytridiomycosis, caused by the micro-fungus *Batrachochytrium dendrobatidis* (*Bd*). Previously, it has been shown that several micrograzers may consume *Bd* zoospores [12]. We expand on this by first indicating that some protozoa can also grow on *Bd* zoospores for short periods, and that the ubiquitous ciliate *Tetrahymena pyriformis* grows at near maximal rates on zoospores alone for extended periods. Recognising that micrograzers will grow on *Bd*, provides essential information for modelling population dynamics. We then provide an assessment of the functional response of *T. pyriformis* feeding on spore*-*sized prey, also required for establishing a population model. Finally, using our data and literature estimates, we develop and employ a novel model that couples the *Bd* life cycle with micrograzer-control on zoospores. This synthesis reveals the dynamics by which micrograzers may supress *Bd* loads and argues that for theoretical predictions of *Bd*-host interactions it will be useful to consider embedding these into the larger microbial food web.

### *Micrograzer growth on* Bd

Previous work has suggested that a large range of micrograzers will consume *Bd* zoospores and may reduce *Bd* viability[12]. We have extended this observation by assessing if *Bd* provides nutritional benefits, that support micrograzer growth. All of the micrograzers we examined could ingest *Bd*, but they exhibited a range of growth-responses. For one ciliate, *S. coeruleus, Bd* appeared to be toxic (possibly explaining previous reports that *S. coeruleus* does not reduce *Bd* viability;[12]), while other species seemed to obtain no nutritional benefit (Fig. 2). However, several species benefited from ingesting *Bd*. Both *Paramecium aurelia* and *P. caudatum* initially grew, although this was not sustained (Fig. 3), suggesting that while *Bd* is of some value, it may lack essential nutrients for these ciliates. In contrast, *Tetrahymena pyriformis* sustained positive growth, indicating that *Bd* can provide a complete diet for certain species. These observations are supported by previous work on ciliates: *T. pyriformis* and a closely related species, *Colpidium striatum*, also grow on yeast (*Saccharomyces*), while *P. aurelia*, and *P. caudatum* cannot, again possibly due to a lack of nutrients such as essential fatty acids and B-vitamins [38, 39].

Our analysis, therefore, suggest that not all micrograzers would be capable of or equally proficient at controlling *Bd*. However, with additional prey sources to sustain the consumers, there may be selective feeding on *Bd*. For instance, *T. pyriformis* differentiates between prey, leading to a more efficient assimilation of prey biomass and a greater cell yield of ciliates [40]. Likely, in the mesocosm experiments conducted by Schmeller et al. [12], where *Paramecium* controlled *Bd*, this ciliate’s diet was supplemented by naturally occurring bacteria. In fact, in our initial growth-experiments, where antibiotics were not included, bacteria grew, and *Paramecium caudatum* consumed zoospores in addition to bacteria and maintained extended positive growth (Supplement 1). Our analysis here has focused on *Bd* as the sole food source and indicates that micrograzer dynamics (growth and ingestion) lead to control of *Bd* populations. These findings argue that *Bd*-host dynamics should now be examined in a wider food-web context, with mixed an assemblage of microgazers sustained by *Bd* and a wider range of natural food sources.

### *Tetrahymena grazing on* Bd

Globally, *Tetrahymena* is common in shallow waters, living near sediments, where it consumes bacteria and other microbes [41, 42]. These are the same habitats that *Bd* occupies. *Tetrahymena* is also associated with amphibians where it may be an opportunistic ectoparasite [43, 44], but possibly also a consumer of *Bd* zoospores as they emerge from sporangia. Considering its habitat and ability to rapidly reproduce on *Bd* alone, we focused on *T. pyriformis’* ingestion of *Bd* zoospores. In contradiction to Schmeller et al., attempts to stain *Bd* zoospores with calcofluor-white [12] were not successful; calcofluor stains chitin [45], and although *Bd* sporangia have a chitin wall, zoospores do not [28]. Therefore, this staining method seems inappropriate for *Bd* zoospores. We then explored vital stains (e.g. cell tracker green), but again we were not successful. Consequently, ingestion estimates relied on the uptake of fluorescent beads as surrogates for *Bd*, which may underestimate rates (e.g. [46]). We, therefore, see our predictions as conservative. From our data, a clear Type II functional response was obtained for *T. pyriformis* (Fig. 4), providing essential parameters for modelling *Bd*-micrograzer dynamics (see Methods). To our knowledge, this is the first time a functional response on *Bd* sized particles has been obtained for any *Tetrahymena* species: the estimates of *I*_max_ and *k* are within the range of those obtained for other ciliates, although the *k*-values are on the lower end of the spectrum [39-47, 48]; our modelling, therefore, includes micrograzers that are inferior to *Tetrahymena*.

### *Modelling micrograzer impacts on* Bd *populations*

Empirical evidence suggests that *Paramecium* can reduce *Bd* infections, through examining end point estimates of host infection [12]. Here we explore the temporal dynamics of such control and the potential for micrograzers to prevent host death. Our analysis is reductionist and hence more qualitative than quantitative in its predictions. However, it clearly reveals that by applying plausible parameters for both the parasite and micrograzer, in a local environment, chytridiomycosis may at times be prevented and *Bd* virtually eradicated, or at least reduced to negligible levels (Fig. 6). Critically, it suggests the time scales over which such dynamics may occur. Admittedly, we indicate that micrograzers that are inferior to *T. pyriformis* are less likely to prevent host death, yet they still, ultimately, reduce *Bd* populations to negligible numbers, potentially preventing further disease outbreaks (Fig, 6). Our model, therefore, provides a mechanism to evaluate *Bd*-micrograzer dynamics, and its predictions strongly argue for the continued exploration of micrograzers in *Bd* research, specifically, and in the control of a range of diseases that spread through zoospores or other similarly sized dispersal stages [3-5].

To date, models of *Bd*-dynamics [8, 9, 49, 50] have not included estimates of spore loss by micrograzers. As indicated above, the modelling provided here is instructive and could benefit from elaboration. Given the ubiquity of protozoa in natural waters [2], and the clear indication of their potential impact (Fig. 6, [3-5]), we suggest there is now a need for better parameterization of micrograzer-*Bd* responses. We suggest three main directions. First, micrograzers, and specifically *Tetrahymena*, exhibit chemosensory behaviour [35]; the extent to which protozoa are attracted to amphibian hosts and *Bd* requires evaluation. Second, as indicated above, the role of *Bd* as a supplement rather than a sole dietary component deserves attention. Finally, the Rosenzweig-MacArthur predator-prey model, which we used, has limitations. Model structures such as the independent response model [51] that rely, independently, on growth and ingestion responses provide better predictions [52]. To this end, we suggest that both functional (ingestion) and numerical (growth) responses associated with *Bd* abundance are established for a range of micrograzers.

### *Future directions for microbial ecology and* Bd

Both *Tetrahymena* and *Paramecium* are common species in shallow waters [41-42, 53], that, as we have shown, are capable of growth on *Bd* zoospores for limited to extended times. Undoubtedly, other protozoa will also consume and grow on *Bd* zoospores. We, therefore, suggest that the role of micrograzers is considered when evaluating *Bd*-dynamics and the dynamics of other zoosporic diseases. Contributions of micrograzers to disease dynamics are also likely to have a highly site-specific component, due to their dependence on environmental factors [12]. For instance, chytridiomycosis is more prevalent at higher altitudes [34], which will often be both cooler and oligotrophic. While temperature may, in part, determine infection burdens [54], there will likely be an interaction with the trophic status of the water. If in oligotrophic waters bacteria are reduced below levels sufficient to support ciliates (<10^6^ ml^-1^), top-down control may be absent, and our analysis suggests that *Bd* may thrive, resulting in chytridiomycosis. Consequently, assessing the abundance of micrograzers in waters where chytridiomycosis occurs or is predicted seems warranted.

We end with some speculations on the potential for biomanipulation using micrograzers, as this has been considered by others [12]. Traditional approaches for managing wildlife diseases have proven ineffectual or impractical for *Bd*. Treating amphibians with probiotic bacteria directly has generally been unsuccessful as a conservation strategy (but see [34]), and, while theoretically plausible, culling hosts to below the critical threshold for disease transmission contravenes conservation objectives [55-57]. This means that alternative mitigation and management strategies must be perused. To date, the only successful effort towards in situ *Bd*-mitigation relied on dosing animals with antifungal chemicals alongside applying chemical disinfectants directly to the environment to reduce spore survival [55]. Fungicides and chemical disinfection, however, have shortcomings, not least of which are ethical issues associated with indiscriminate toxic effects. As both *Tetrahymena* and *Paramecium* are already common if not ubiquitous in aquatic environments and are simple and inexpensive to grow in large quantities [42, 53], they may be tractable target species for biomanipulation. We, therefore, support previous suggestions Schmeller et al. [12] that by augmenting natural densities of these species, through addition or supplementary feeding, it may be possible to reduce zoospore densities for *Bd* in situ. However, an introduction or population increase of any species could have detrimental, ecosystem-changing effects that require in depth evaluation before application [58]. Further evaluation of the role of microgazers is, therefore, required before we can understand their likely impact in natural conditions, and advocate the implementation of such approaches.

## Acknowledgments

This work was conducted as part of postgraduate project (HF) at the University of Liverpool and was supported by funding provided to JJ through the Program for Training University Teachers by Shanghai Municipal Education Commission. AF and TG were funded by the NERC grant NE/N009800/1. MCF is a CIFAR fellow with *Bd* research supported by NERC NE/K014455/1. Eleanor Ambrose, Philip Bahan, Valentina Iorio, and Sam Moss provided invaluable technical support.

## Conflict of Interest

The authors declare that the research was conducted in the absence of any commercial or financial relationships that could be construed as a potential conflict of interest.

## Notes

### Competing Interest Statement

The authors have declared no competing interest.

